# Bridging the neural and algorithmic correlates of action-stopping

**DOI:** 10.1101/2022.10.06.511081

**Authors:** Sumitash Jana

## Abstract

Rapid stopping of actions is crucial for survival. The algorithm underlying action-stopping is often described using the independent race model, where the race between independent go- and stop-processes determines whether one is able to stop. However, it is unclear whether humans have neural correlates of these processes. Here, using scalp potentials from three studies, we show that the mu (8-13 Hz) desynchronization over the contralateral motor cortex follows the predictions of a go-process: the activity builds up till an action is made. We also show that the low beta (13-20 Hz) activity over the frontal areas follows the predictions of a stop-process: the activity builds up after the stop signal and prior to the stopping latency. Using these neural correlates, we observed that: 1) The failure to stop mostly results from a rapid go-process, 2) The go-process seems to be influenced by the stop-process during the late stages of action-stopping, 3) Intrinsic prefrontal beta activity has a retardive effect on movement, 4) Violations of independence assumption in the failed stop trials may result from the activity of the stop-process. Future studies may use these neural correlates to resolve the mimicry between models of action-stopping.

## Introduction

In our constantly changing environment, we must rapidly control our actions when the need arises. For example, while driving when the traffic lights turn red we must stop pressing the gas pedal and press the brakes instead. While such form of control probably requires many cognitive processes, one critical process is action-inhibition, the cognitive process that mediates the behavior of rapid stopping of actions (action-stopping). In the lab, action-stopping has been extensively studied using the stop-signal task (Verbruggen et al., 2019). In this task, participants make an action in every trial, but in a minority of ‘stop trials’ when a stop-signal is presented, they must try to stop the incipient action. Our comprehension of action-stopping has significantly progressed at all three levels of understanding – the computation, algorithm, and the implementation levels (Marr, 1982), but these investigation have largely remained confined within each levels. At the algorithm level, numerous models have been proposed to how one is able to successfully stop or fail to stop in a stop trial (Bissett et al., 2021; Boucher et al., 2007; Indrajeet and Ray, 2019; Logan et al., 2015; E. Salinas and Stanford, 2013). Probably, the simplest and most popular version is the independent race model which suggests that the outcome of a stop trial is determined by a race-to-threshold between a go-process and a stop-process (Logan and Cowan, 1984). The prokinetic go-process starts when the go cue is presented and the akinetic stop-process starts when the stop-signal is presented. If the go-process reaches the threshold first then one fails to stop the action, and if the stop-process wins the race then one successfully stops the action. Despite several criticisms of this model (Bissett et al., 2021; Boucher et al., 2007; Colonius and Diederich, 2018; Logan et al., 2015), it still remains popular partly because this model also provides a way to estimate the latent time taken to stop a response called Stop Signal Reaction Time (SSRT), a measurement that cannot be directly made from behavior. At the implementation level, a prefrontal-basal ganglia-thalamocortical pathway is thought to be important for stopping actions (reviewed in (Hannah and Aron, 2021; Wessel and Aron, 2017)) (also see (Diesburg and Wessel, 2021)). However, the link between the algorithm and implementation levels of action-stopping is unclear. In other words, despite the widespread use of the independent race model (and its variations), it is worth noting that it is not established whether the race model has neural correlates in humans or whether it just provides a theoretical framework for understanding action-stopping. It must be noted that while evidence is lacking in humans, neural correlates of the go- and stop-processes have been proposed in rodents and non-human primates: The temporal dynamics of certain neural populations in the basal ganglia in rats (Schmidt et al., 2013) and in the frontal eye fields in monkeys (Hanes et al., 1998) resemble the activity of the go- and stop-process. However, there are several caveats to drawing inferences from these studies: these studies were performed in a small sample of animals, the animals were highly over-trained, only small populations of neurons were sampled, and lastly but most importantly, it unknown whether these findings generalize to humans. The current study is an attempt to bridge this gap by testing the existence of neural (electroencephalography, EEG) correlates of the go- and stop-processes in humans.

Establishing the neural correlates of the go- and stop-processes can have several benefits. First, it will help us study the temporal dynamics of action-stopping and might help resolve extant questions such as: Are stop failures just dependent on the race between the go- and the stop-process, or does it also depend on the failure to ‘trigger’ the stop-process (Band et al., 2003; Jana and Aron, 2021; Logan and Cowan, 1984; Matzke et al., 2017)? Second, it will allow us to study the temporal dynamics of action-stopping in clinical populations who suffer from inhibitory control deficits which might lead to development of better therapeutic interventions. Third, it might help development of better models to understand action-stopping. As mentioned above, there are numerous other models that can explain action-stopping such as interactive race model (Boucher et al., 2007; Logan et al., 2015), deceleration model (Indrajeet and Ray, 2019; Emilio Salinas and Stanford, 2013), and blocked-input model (Logan et al., 2015). Many of these models have similar performance and it is difficult to break the model mimicry by computational studies alone. Relatedly, (Bissett et al., 2021) has recently demonstrated that there are violations of assumptions of the independent race model at short stop signal delays (SSDs) and has recommended the development of newer models. Thus, studying the neural correlates of the go- and stop-process can help break model mimicry (Logan et al., 2015) and can inform the development of better models. Fourth, these neural correlates might extend to inhibitory control in other domains such as inhibition of urges and thoughts or action-stopping in real-world scenarios and thus galvanize the understanding in these domains.

Now, which EEG signals could relate to the go- and stop-process? One obvious target for the neural correlate of the go-process would be the activity recorded over primary motor cortex during manual actions. Previous research has shown that during movement preparation, mu/alpha (8-15 Hz) desynchronization is observed over the contralateral motor cortex (Crone et al., 1998; Engel and Fries, 2010; Pfurtscheller and Lopes da Silva, 1999). Interestingly, other studies have shown that there is greater desynchronization in the failed stop compared to successful stop trials (Fonken et al., 2016; Swann et al., 2009) which might indicate a faster go-process in the failed stop trials. Hence, we chose the mu/alpha desynchronization as a potential correlate of the go-process. Such a suggestion has been made previously in a decision task (O’Connell et al., 2012). To converge on a potential neural correlate of the stop-process, we used Independent Component Analysis (ICA) to isolate the low beta activity (13-20 Hz) over right frontal EEG electrodes that is thought to be involved in action-stopping (Castiglione et al., 2019; Hannah et al., 2020; Jana et al., 2020; Wagner et al., 2018). Furthermore, the beta activity of this independent component (IC) is disrupted by repetitive transcranial magnetic stimulation to right inferior frontal gyrus (Sundby et al., 2020), which is a key node in the prefrontal-basal ganglia-thalamocortical network that mediates action-stopping (reviewed in (Hannah and Aron, 2021; Wessel and Aron, 2017)). Beta power in this IC is greater during successful stopping (Sundby et al., 2020), and beta bursts in this IC is correlated with muscle index of stopping, even on a single trial level (Hannah et al., 2020). We decided against using beta bursts because they occur in ~15% of trials (Errington et al., 2020; Jana et al., 2020) indicating that may be neither necessary nor sufficient for stopping (Errington et al., 2020; Muralidharan et al., 2022). Hence, we chose the time-course of beta power in the right prefrontal EEG component as a potential correlate of the stop-process.

In three existing human EEG data sets, we first tested whether our proposed neural correlates of the go- and stop-process followed the predictions of the race model. We then tried to address several questions regarding action-stopping behavior and the race model: 1) What determines stop failures, a faster go- or slower stop-process? 2) Is the go-process independent? 3) Do behavioral context violations as demonstrated by (Bissett et al., 2021) result from a weak stop-process? 4) What is effect of frontal beta in go trials?

## Results

Behavioral performance in the stop-signal task was typical in all three data sets (***Table 1***). Accuracy in go trials was >98% with roughly 50% successful stopping in stop trials. SSRT was also typical with the mean across the data sets ranging between 219 - 232 ms. In all three data sets, the mean failed stop RT was lesser than the mean correct go RT.

**Table 1:** Behavior (mean±s.e.m)

|  | Data set 1 | Data set 2 | Data set 3 |
| --- | --- | --- | --- |
| **Correct Go RT (ms)** | 463±16 | 407±6 | 414±8 |
| **Failed Stop RT (ms)** | 406±14 | 372±6 | 375±6 |
| **Correct Go (%)** | 98±0 | 99±0 | 99±0 |
| **Correct Stop (%)** | 51±1 | 50±0 | 47±1 |
| **Mean SSD (ms)** | 221±15 | 173±6 | 171±10 |
| **SSRT (ms)** | 223±6 | 219±7 | 232±6 |

### Neural correlate of the go-process

Since mu/alpha desynchronization in motor cortex is seen during movements (Crone et al., 1998; Engel and Fries, 2010; Pfurtscheller and Lopes da Silva, 1999), we started by testing whether the mu/alpha power in the electrodes over the contralateral motor cortex (electrodes C1, C3, C5; left hemisphere, LH) followed the predictions of the go-process (see *Methods*; ***Figure 1 – figure supplement 1A***). We reasoned that a neural correlate of the go-process should satisfy two criteria: 1) It should show build-up of activity with time, starting with the presentation of the go cue and reaching a peak at the time of the response, 2) The activity should vary with reaction time (RT) such that the activity in slower response trials should have a slower rate of build-up.

The mu/alpha desynchronization satisfied these criteria. We observed that it started ~100 ms following the go cue and increased till the response was executed. This rise-to-peak activity at or before the response, was more evident when the activity was aligned to RT, thus satisfying criterion 1 (***Figure 1 – figure supplement 1B-C***). Next, we tested whether there was a RT-related modulation in activity. To test this, we divided the go RT distribution into faster and slower halves using the sum of the participant’s mean SSD and SSRT (Schmidt et al., 2013). The activity was faster and/or stronger in the fast vs. slow RT trials, thus, satisfying criterion 2. When aligned to the time of response, the activity in the slow RT trials started earlier than that in the fast RT trials in all data sets (analysis period starting from mean time of go cue till response) (***Figure 1A-C***). We reasoned that this difference in RT might be related to intrinsic neural variability or might be related an inhibitory influence. We consider the latter possibility below. Taken together, this indicates that the mu/alpha desynchronization over the contralateral motor cortex may be considered as the putative go-process (called go-related activity henceforth).

**Figure 1.**
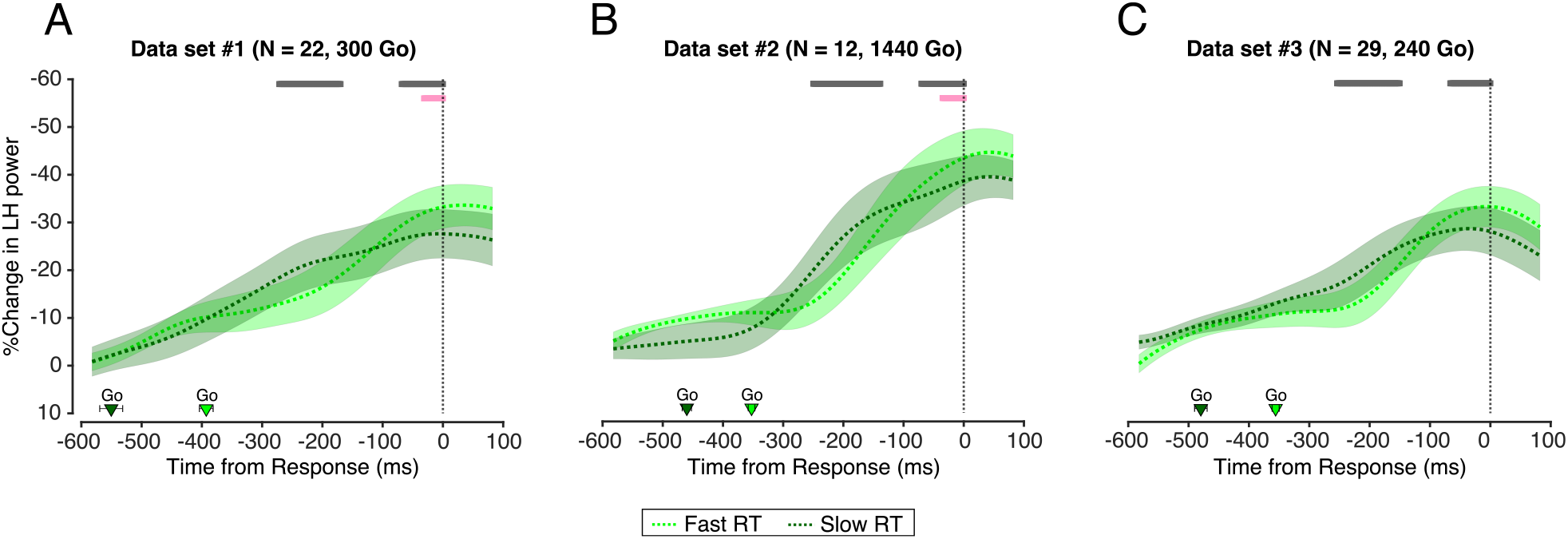
Neural correlate of the go-process. **A-C**) Percent change in the left hemisphere mu/alpha power with respect to baseline (−400 to 0 ms prior to go cue) aligned to the time of response in data set 1-3, respectively. The dotted line and shaded region indicate the mean±s.e.m. across participants in data set 1 (light green – fast RT, dark green – slow RT). The triangles and cross-hairs at the bottom indicate the mean±s.e.m. of the time of presentation of the go cue across participants. The pink and grey bars at the top indicate time-points where the difference between the slow and fast RT group was statistically significant using non-parametric permutation testing and non-parametric bootstrap, respectively.

### Neural correlate of the stop-process

Since low beta power (13-20 Hz) over right frontal brain areas has been implicated in action-stopping (Castiglione et al., 2019; Sundby et al., 2020; Swann et al., 2009; Wagner et al., 2018), we tested whether the dynamics of low beta activity over right frontal electrodes followed the predictions of the stop-process (see *Methods*). To identify electrodes of interest in each participant (i.e. a spatial filter), we used Independent Components Analysis (Bell and Sejnowski, 1995) (also see (Castiglione et al., 2019; Wagner et al., 2018)) and chose a brain-related independent component (IC) for each participant based on two criteria. First, the scalp topography had to be right-frontal (if it was not present, then frontal), and, second, increased low beta power in the successful stop trials (stop signal to SSRT) compared to the fixation period (Hannah et al., 2020; Jana et al., 2020). Based on the race model, we reasoned that a neural correlate of the stop-process should satisfy three criteria, 1) It should show greater-than-baseline build-up of activity following the presentation of the stop signal and reach a peak before or at SSRT in stop trials, 2) Its activity should be greater in stop compared to go trials, 3) Its activity should increase earlier at shorter SSDs compared to longer SSDs.

The right frontal beta activity satisfied the criteria of a putative stop-process. The low beta activity increased after the stop signal and reached a peak at or before SSRT in the stop trials thus satisfying criterion 1 (***Figure 2A-C***). Further, this activity was significantly greater in the stop trials compared to go trials particularly before SSRT, thus satisfying criterion 2 (***Figure 2A-C***) (***Figure 2 – figure supplement 1A-F*** for the comparison between successful stop vs. go trials). In all data sets, this difference appeared 50-100 ms after the stop signal and lasted till SSRT. This build-up was observed in the successful stop trials which had greater activity than slow go RT trials. Note that the activity also build-up in the go trials, albeit lower than that in the stop trials. We consider this build-up in the go trials later. As a control, we tested for the specificity of both the frequency and location of the IC. First, the activity of high beta (21-30 Hz) in the same ICs often did not increase beyond the baseline in the stop trials highlighting the frequency-specificity (***Figure 2 – figure supplement 2A-C***). We chose high beta because the entire beta band (13-30 Hz) has often been implicated in action-stopping (Aron et al., 2016; Schmidt et al., 2019). Second, low beta activity in the stop trials in a posterior IC did not increase beyond the baseline highlighting the location-specificity (***Figure 2 – figure supplement 2D-F***). We chose the posterior IC as it would not be contaminated by volume conduction and because this IC lies over areas that show significant activity during the stop signal task (Osada et al., 2019) but may not have a direct relevance to action-stopping (Hannah and Jana, 2019).

**Figure 2.**
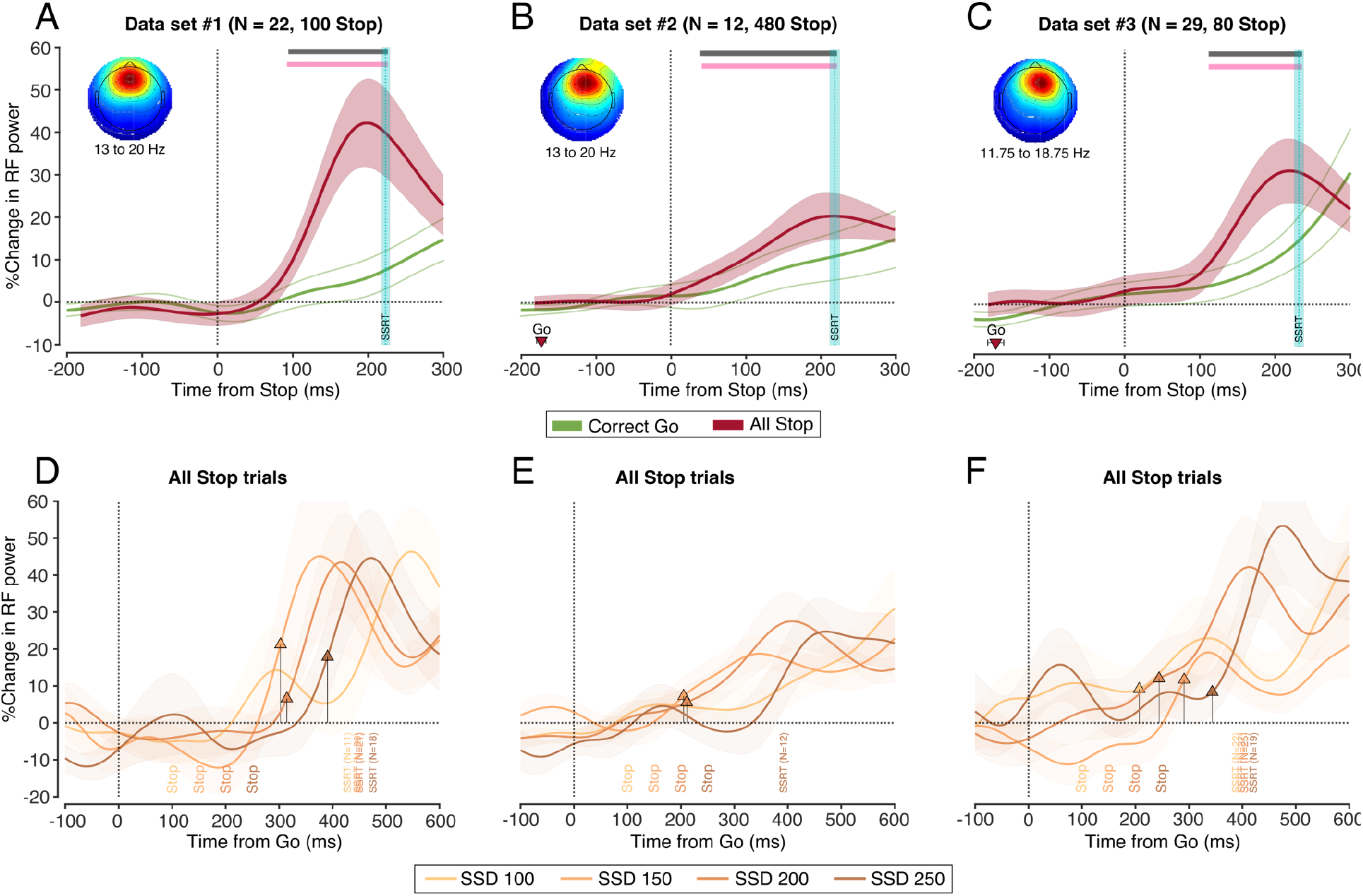
Neural correlate of the stop-process. **A-C)** Percent change in the right frontal low beta power with respect to baseline (−400 to 0 ms prior to go cue) aligned to the stop signal in data set 1-3, respectively. The solid line and bounded region indicate the mean±s.e.m. across participants in data set 1 (red – successful stop, dark green – slow RT go trials). The activity of the go trials is pseudo-aligned to the SSD in the successful stop trials. The triangles and cross-hairs at the bottom indicate the mean±s.e.m. of the time of presentation of the go cue in the successful stop trials across participants (unless it is outside the x-axis range). The dotted cyan line and shaded region represents the mean±s.e.m. of SSRT across participants. The pink and grey bars at the top depict time-points of statistical significance using non-parametric permutation testing and non-parametric bootstrap, respectively. **D-F)** Percent change in the right frontal low beta power with respect to baseline (−400 to 0 ms prior to go cue) in the most common SSDs (100, 150, 200, and 250 ms) aligned to the go cue in data set 1-3, respectively. Each SSD is represented as text ‘Stop’ at the bottom. The arrow marks the first time-point after the SSD when the RF power increases above 0 (see *Methods*). The SSRT across participants who contributed to each SSD is marked as text at the bottom of each figure.

Finally, we tested whether this activity had a parametric relationship with SSD. To test this, we aligned the low beta activity at each SSD to the go cue (see *Methods* for details) and observed the time when the activity increased from baseline after the stop signal. In all data sets and most SSDs, the parametric relationship was generally maintained where the activity increased earlier for short compared to long SSDs (***Figure 2D-F***). In data set 1, the relationship was evident where the activity of SSD 150 ms increased ahead of SSD 200 ms which increased ahead of SSD 250 ms. In data set 2, a similar pattern was observed. However, in data set 3, the activity for SSD 200 ms appeared before 150 ms, but the parametric relationship was seen for SSDs 100, 150, and 250 ms. Note that for some SSDs we were not able to detect a significant increase in activity prior to SSRT. This might be because there were few trials for some SSDs (cutoff of 10 trials per SSD per participant) and because of noise that is inherent to EEG. Nevertheless, the fact that a similar pattern was observed in all the three data sets suggests that it might be a true effect. Thus, we considered that criterion 3 was also satisfied. Taken together, this indicates that the low beta power over the right frontal electrodes may be considered as the putative stop-process. We call this stop-related activity henceforth.

Thus, having proposed the putative go- and stop-process, we then tested whether the dynamics of these processes could inform about several extant questions of action-stopping.

### Why does one fail to stop in some trials?

Next, using the go-related and stop-related activities we tracked the relationship between the two processes in stop trials. Specifically, we tried to answer a basic question of stop signal behavior: why does one fail to stop in some trials? According to the race model, stop failures can result from either a fast go-process or a slow stop-process. A fast go-process would mean that in failed stop trials, go-related activity should be greater than that in the successful stop trials, especially early on in the trial. A slow stop-process would mean that in failed stop trials, stop-related activity would be less than that in the successful stop trials, especially prior to SSRT. To test this, we binned the time from the stop signal into 50 ms bins, and tested the difference in the go-related and stop-related activity between the failed and successful stop trials (***Figure 3A-C***). A fast go-process seemed to be the main determinant of stop failures. In all 3 studies, the go-related activity was greater in failed stop compared to successful stop trials especially early in the trials. This difference was visible even prior to the stop signal and often persisted till the mean failed stop RT. The contribution of the stop-related activity in determining the outcome of a stop trial was lower than the contribution of the go-related activity as there was often no significant difference in stop-related activity prior to SSRT. This result was invariant to the bin duration (data not shown). Taken together, this validates the theoretical account of the race model, where a go and stop-process race towards threshold, and the relative time when they reach the threshold determines the outcome of the trial. Our results suggest that one fails stop in some trials because the go-process reaches the threshold before the stop-process can intervene.

**Figure 3.**
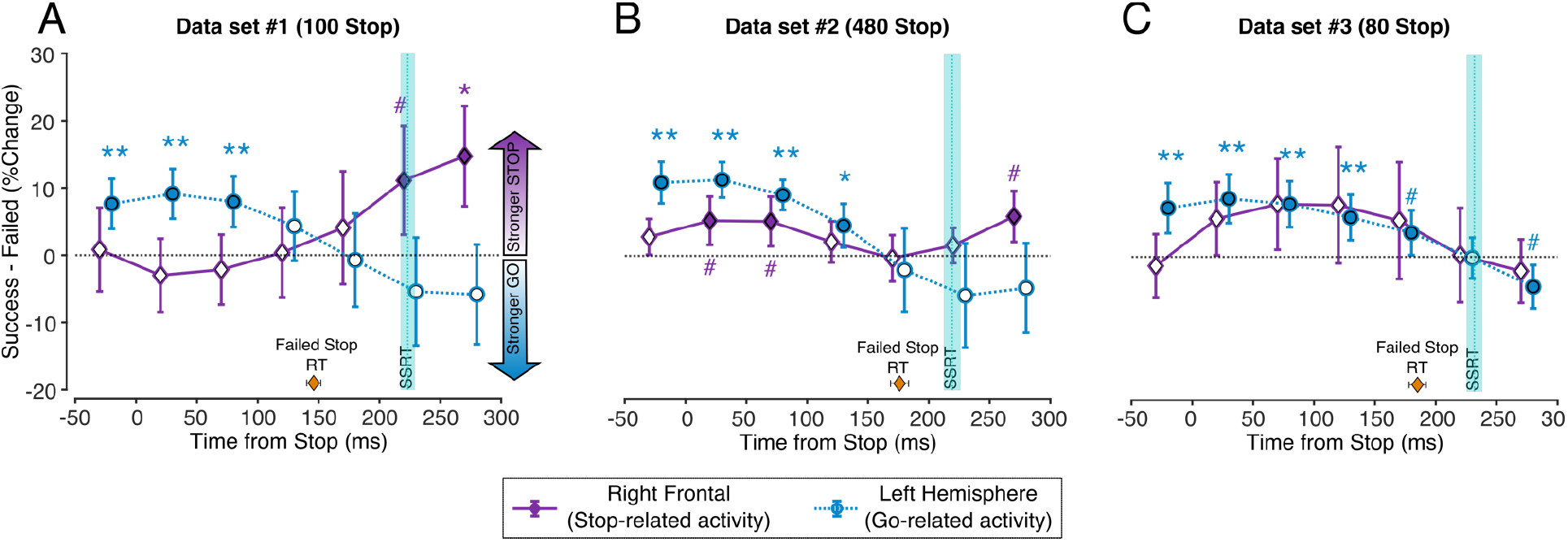
Relationship between go-related and stop-related activity in stop trials. **A-C)** The go-related and stop-related activity represented as the activity difference between the successful and failed stop trials aligned to the stop signal in data set 1-3, respectively. Each circle and cross-hair represent the mean±s.e.m. across participants in that time-bin (blue – go-related activity, purple – stop-related activity). Filled dots represent statistical significance. The dotted cyan line and shaded region represents the mean±s.e.m. of SSRT across participants. Positive values for the difference in stop-related that it is stronger, and positive values for the go-related activity indicates that it is weaker in successful compared to failed stop in that time-bin, respectively. The diamond shape and cross-hairs at the bottom represents the mean failed stop RT. (^#^, *p*_*BH*_ < 0.1 & *p*_*BH*_ >=0.05; *, *p*_*BH*_ < 0.05 & *p*_*BH*_ >=0.01; **, *p*_*BH*_ < 0.01 & *p*_*BH*_ >=0.001; ***, *p*_*BH*_ < 0.001).

### Is the go-process independent?

Having validated the race between the go- and stop-process in the stop trials, we then tested a key assumption of the independent race model - the independence between the two processes. This assumption allows one to use the go RT distribution as an estimate of RT distribution in the stop trials and thereby allows SSRT estimation. Many studies have questioned this independence assumption based on behavioral, computational, and neural evidence (Bissett et al., 2021; Boucher et al., 2007; Logan et al., 2015). However, some computational studies have clarified that the dependence is seen only in the late stages of the trial, and thus may have a minor effect of SSRT estimations (Boucher et al., 2007; Colonius and Diederich, 2018). Here, we tested this independence assumption by testing how the go-related activity in failed stop trials compared with that in the RT-matched go trials. We reasoned that if the go-related activity was comparable between these two trial types then it would indicate independence of the go-process, else dependence. Across all data sets, when aligned to the time of response, the go-related activity in the failed stop trials diverged from that in the RT-matched go trials ~100 ms prior to the response (***Figure 4A-C***). This suggests that the go-process is not independent and it is consistent with previous computational studies that the dependence is observed only in the late stage of the trial (Boucher et al., 2007; Colonius and Diederich, 2018).

**Figure 4.**
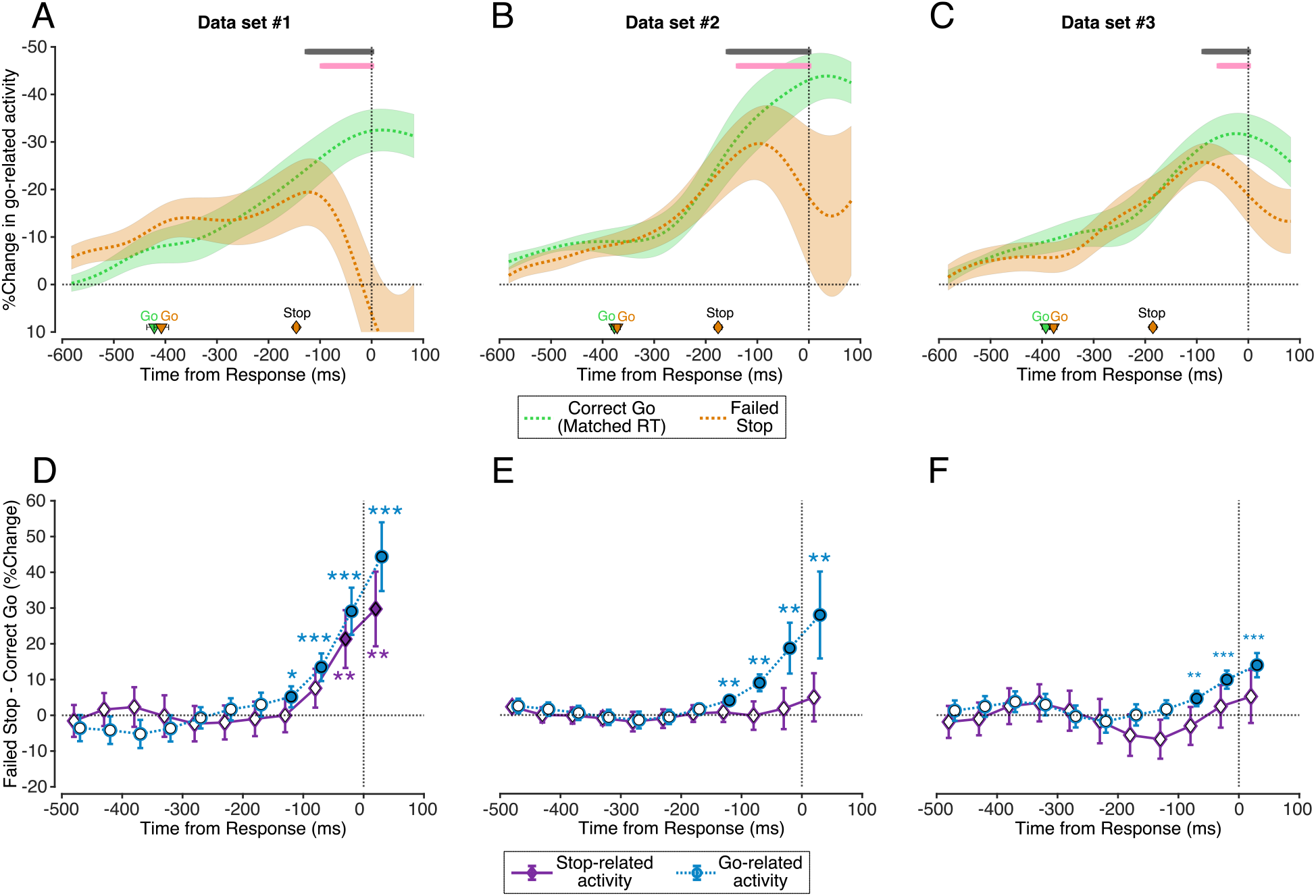
Dependence of the go-related activity. **A-C)** Percent change in the left hemisphere mu/alpha power with respect to baseline (−400 to 0 ms prior to go cue) aligned to the response in data set 1-3, respectively. The dotted line and shaded region indicate the mean±s.e.m. across participants (orange – failed stop, green – latency-matched go trials). The triangles and cross-hairs at the bottom indicate the mean±s.e.m. of the time of presentation of the go cue across participants. The pink and grey bars at the top depict time-points of statistical significance using non-parametric permutation testing and non-parametric bootstrap, respectively. **D)** Difference in the go-related and stop-related activity between the failed stop and latency-matched go trials aligned to the time of response. Each circle and cross-hair represent the mean±s.e.m. across participants in that time-bin (blue – go-related activity, purple – stop-related activity). Filled dots represent statistical significance. Positive value for the difference in stop-related and go-related activity indicates that stop-related activity is stronger and go-related activity is weaker in successful compared to failed stop in that time-bin, respectively. (*, p_BH_ < 0.05 & p_BH_ >=0.01; **, p_BH_ < 0.01 & p_BH_ >=0.001; ***, p_BH_ < 0.001). **E-F**) Same as D but for data sets 2-3.

Now the question arises, whether the change seen in the go-related activity is due to the influence of the stop-related activity. It is possible that in the failed stop trials, the stop-related activity intervenes at a late stage and reduces the go-related activity but this intervention is too late to change the outcome of the trial (i.e. a ballistic stage, discussed below). To test this, we again binned the difference in activity in the go-related and stop-related between the failed stop and RT-matched go trials (***Figure 4D-F***). We reasoned that if the two are related then the time-course of the change in go-related activity should relate to the time-course of change in the stop-related activity. In data set 1, the stop-related activity in the failed stop trials showed a rapid increase prior to the response, and the change in stop-related activity was similar to the change in the go-related activity. However, this was unclear in the other two data sets. One possibility is that the magnitude of change in sensorimotor areas is just typically higher than that in the frontal areas. Another possibility is that the strength of inhibition is greater in stop trials. In other words, in stop trials, there is an executive recruitment of prefrontal beta in stop trials (over and above the intrinsic beta, see next section) and a small change in beta activity can result in a large effect on the go-related activity. In computational models, this is usually parameterized as the interaction term that modulates the ‘strength’ of inhibition. Taken together, this might indicate that the stop-related activity in the failed stop trials intervenes late in the trial and reduces go-related activity.

### What is influence of the stop-related activity in the go trials?

Recent studies have demonstrated that intrinsic beta activity (bursts) can have a retardive effect on behavior and reduce corticospinal excitability (Hussain et al., 2019; Little et al., 2019; Shin et al., 2017). Here, we hypothesized that the intrinsic the stop-related activity might have a retardive effect on the go-related activity and behavior (note that there is above baseline stop-related activity even in go trials, ***Figure 2A-C***). To test this, we checked whether stop-related activity was greater during in slow RT trials compared to fast RT trials. Indeed, this was the case. The stop-related activity was greater in the slow RT vs. fast RT trials for a considerable duration during the trial in 2/3 data sets and just prior to the response in one data set (***Figure 5A-C***). This increased intrinsic beta activity in the slow RT go trials might highlight the inhibitory influence of prefrontal low beta.

**Figure 5.**
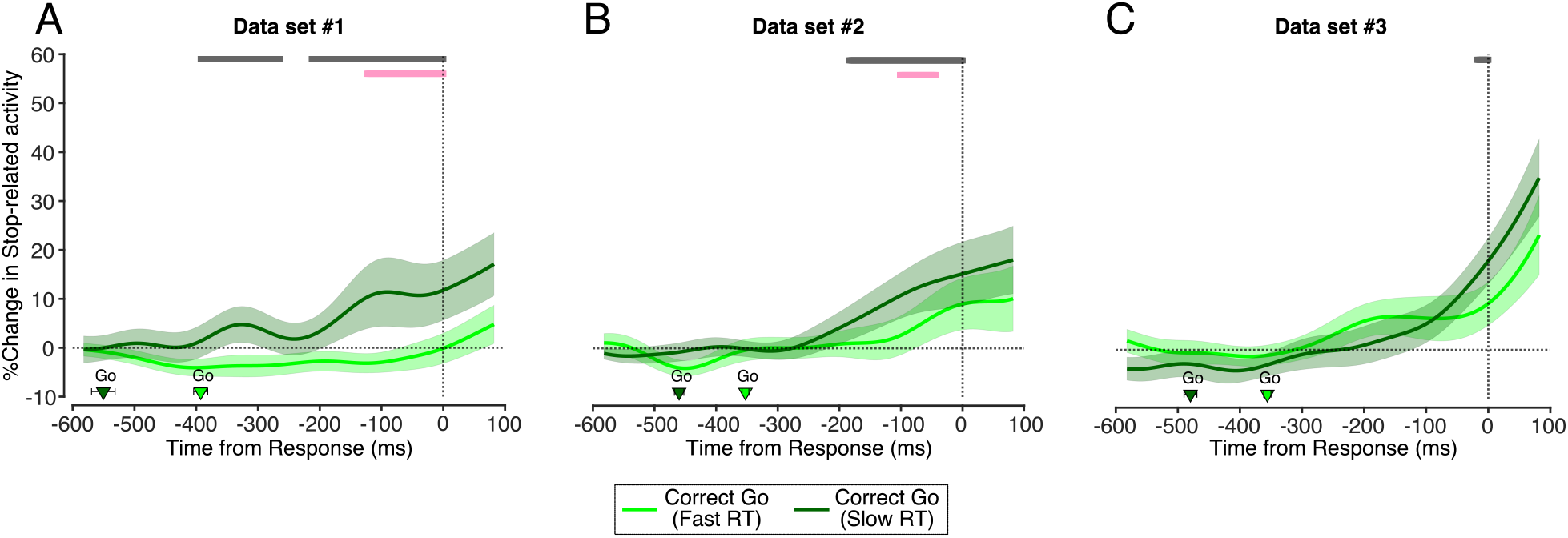
Role of intrinsic beta in go trials. **A-C)** Percent change in the right frontal low beta power with respect to baseline (−400 to 0 ms prior to go cue) aligned to the time of response in data set 1-3, respectively. The solid line and bounded region indicate the mean±s.e.m. across participants in data set 1 (light green – fast RT, dark green – slow RT). The triangles and cross-hairs at the bottom indicate the mean±s.e.m. of the time of presentation of the go cue across participants. The pink and grey bars at the top depict time-points of statistical significance using non-parametric permutation testing and non-parametric bootstrap, respectively.

### What leads to violations of context independence in failed stop trials?

Next, we tested violations of context independence that has been demonstrated by a recent behavioral study (Bissett et al., 2021). In that study, the authors compared the failed stop RT with the prior go RT, and showed that failed stop RTs were often longer than go RTs, especially at short SSDs. They suggest that such violations might result from a weak stop process that acts on the go-process thereby slowing but not stopping the response. Since at short SSDs the influence of the stop-process lasts longer, violations are more common. We tested this hypothesis about a long-lasting but weak stop-process in violation trials. According to this hypothesis, we predicted that the stop-related activity in the violation trials should begin earlier than that in the SSD-matched non-violation trials.

Behaviorally, on an average, we observed that the mean failed stop RT exceeded the prior go RT only at the shortest or longest SSDs not in the middle SSDs (***Figure 6A-C***). However, although across the sample in the middle SSDs, the mean RT difference was negative, there were some participants who had violations even at those SSDs (see the numbers on ***Figure 6A-C***).

**Figure 6.**
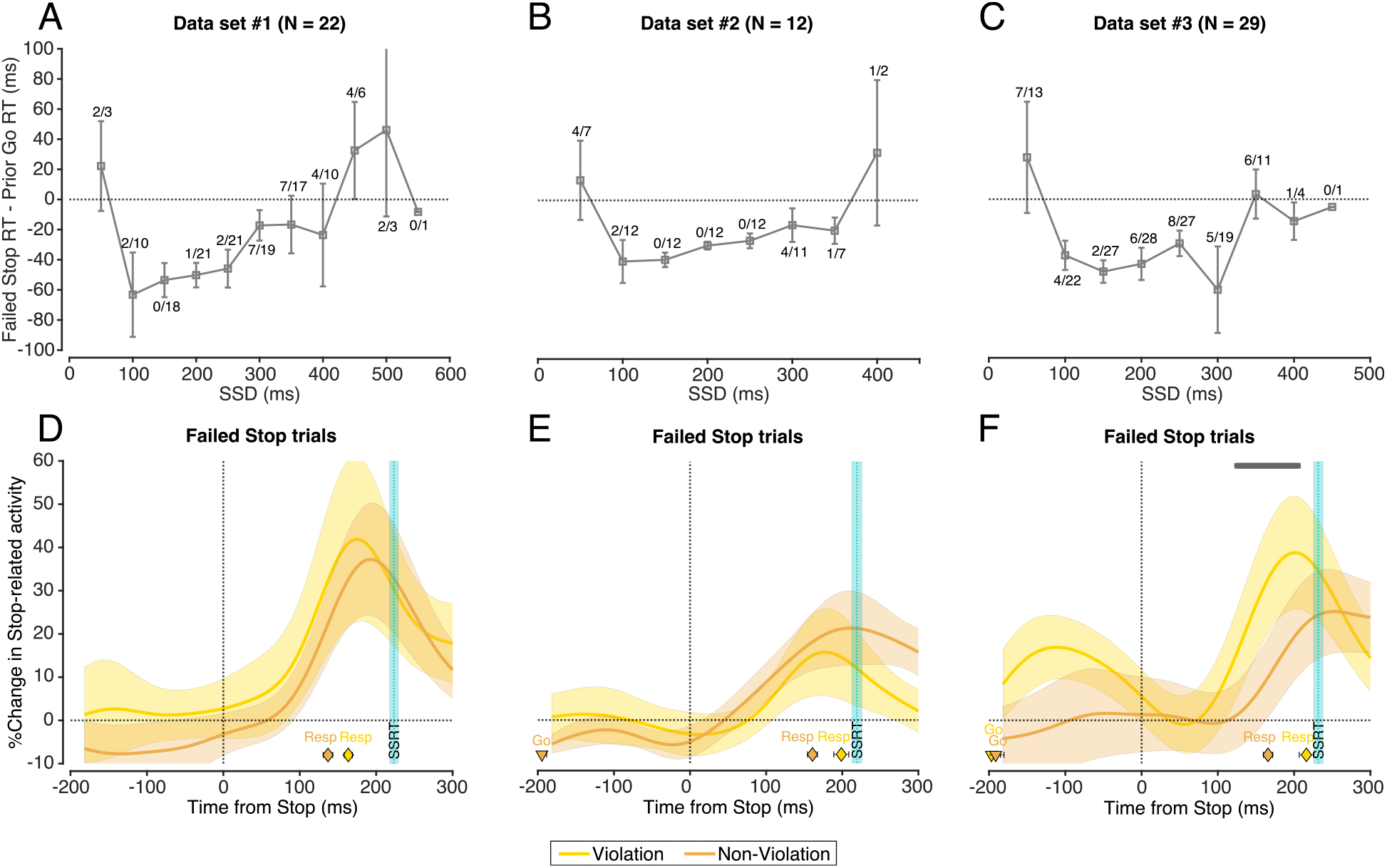
Violations of context independence in failed stop trials. **A-C)** Mean difference in the RT between the failed stop and preceding go trial across the sample in data set 1-3, respectively. The squares and cross-hairs represent the mean±s.e.m. across participants. The numerator of the fraction aligned to each square represents the number of subjects who had violations (positive RT difference) at that SSD, while the denominator represents the total number of subjects who contributed data to that SSD. **D-F)** Percent change in the right frontal low beta power with respect to baseline (−400 to 0 ms prior to go cue) aligned to the stop signal in data set 1-3, respectively. The solid line and bounded region indicate the mean±s.e.m. across participants in data set 1 (yellow – violations, orange – non-violation trials). The triangles and cross-hairs at the bottom indicate the mean±s.e.m. of the time of presentation of the go cue across participants. The diamonds and cross-hairs at the bottom indicate the mean±s.e.m. of the time of response across participants. The dotted cyan line and shaded region represents the mean±s.e.m. of SSRT across participants. The grey bars at the top indicate time-points where the difference between the activity in the two trial types was significant.

**Figure 7.**
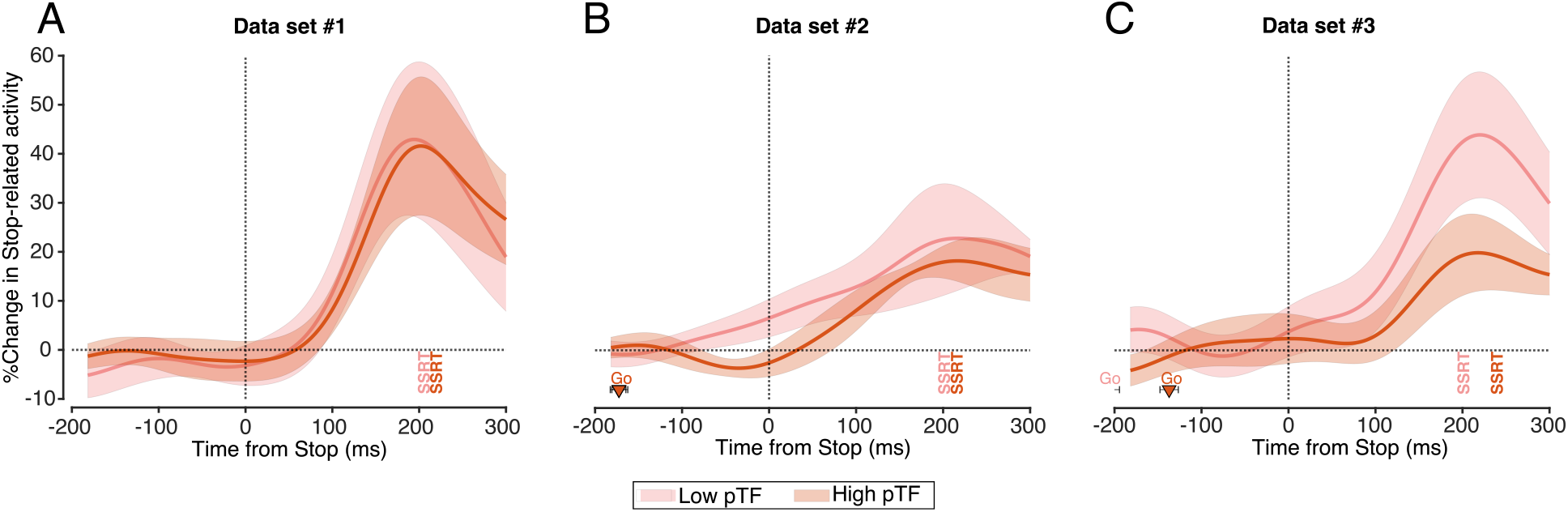
Stop-related activity in participants with low and high probability of trigger failures. **A-C)** Percent change in the right frontal low beta power with respect to baseline (−400 to 0 ms prior to go cue) aligned to the time of response in data set 1-3, respectively. The solid line and bounded region indicate the mean±s.e.m. across participants in study 1 (light red – participants with low probability of trigger failures, dark red – participants with high probability of trigger failures). The triangles and cross-hairs at the bottom indicate the mean±s.e.m. of the time of presentation of the go cue across participants (unless outside the range of the x-axis). SSRT across the sample in each group is also mentioned in the appropriate color.

We then compared the stop-related activity between the SSD-matched violation and non-violation trials. In data set 3, the stop-related activity in the violation trials increased earlier than that in the non-violation trials consistent with the predictions of (Bissett et al., 2021) (***Figure 6D-F***). This preliminary evidence could be further validated in larger data sets to investigate the neural correlates of context violation in some failed stop trials.

### What leads to trigger Failures

Finally, we compared how stop-related activity varied in the stop trials when the probability of trigger failures (pTF) increased. Several recent studies have demonstrated the importance of studying pTF. The probability of trigger failures increases in clinical populations compared to healthy controls (Swick and Ashley, 2020; Weigard et al., 2019), pTF increases when a participant is mind wandering compared to when they are focused on the task (Jana and Aron, 2022), and is modulated by reward (Doekemeijer et al., 2021). A recent study with a large sample size has suggested that the N100 event-related potential amplitude in an auditory stop signal task might be a marker of trigger failures (Skippen et al., 2020). They divided the pTF into bins and showed that N100 amplitude is smaller in those with higher pTF. We reasoned that similarly, participants with greater pTF might have a weaker and/or slower stop-related activity. To test this, we median-split the distribution of pTF across participants and compared the stop-related activity between the low vs. high pTF groups. While there was no significant difference in the stop-related activity at any time-point in any data sets, in data set 3, the low pTF group had the trend of increased stop-related activity which did not reach statistical significance. This needs further validation in larger data sets.

## Discussion

Rapid stopping of actions is a fundamental requirement of life. Here, we investigated the neural correlates of the race model, a model that has been the bedrock of inhibitory control research for the last four decades. We observed that some human EEG signals parallel the theoretical go- and stop-processes, thus bridging the gap between the algorithm that mediates action-stopping and its neural implementation. Further, it validates the race model as an appropriate model to study action-stopping. We show that the temporal dynamics of mu/alpha power over contralateral motor cortex and the low beta power over right frontal electrodes may be considered as a putative go-process and stop-process, respectively. Leveraging the dynamics of these processes, we were able to address several questions. First, we show that stop failures are predominantly related to a fast go-process. Second, we show that the go-process is not independent as the go-related activity in failed stop trials diverged from that in the latency-matched go trials. Further, this divergence in go-related activity without a change in behavior suggests the existence of a ballistic stage that is impervious to the stop-process. Third, we observed preliminary evidence that violations of context independence in failed stop trials might result from the activity of the stop-process. Finally, we also show that intrinsic beta on go trials may have a retardive effect by slowing down RT. Thus, this study links our understanding of action-stopping at the algorithm and implementation levels and may allow experimental verification of other theoretical questions existing in the field.

A key part of our study was visualizing the go-related and stop-related activity that paralleled the theoretical accounts of the go- and stop-process, respectively. This method of data visualization has some benefits. First, this allows us to study the temporal dynamics of the go- and stop-process. Some recent studies have used beta bursts as a marker of the stop-process (Hannah et al., 2020; Jana et al., 2020) as bursts allow for trial-by-trial correlation with muscle activity (Hannah et al., 2020). However, these bursts are rare (occurs in ~15% of trials) and other recent studies have suggested that bursts may not be sufficient to account for stopping behavior (Errington et al., 2020; Muralidharan et al., 2022). Our method of visualization when used in combination with bursts may be more useful in understanding of brain dynamics during action-stopping. Second, this visualization may allow development of better, neurally-inspired models for action-stopping. There are numerous models that can reasonably explain stopping behavior and it is hard to break the model mimicry using simulations (Logan et al., 2015). We suggest that these neural correlates of the go-related and stop-related activity might help resolve such model mimicry. These neural correlates might also allow one to experimentally quantify the interaction term between the go- and stop-process for an interactive race model. Third, it may help in distinguishing the contribution of the go- and stop-process in failures of action-stopping in different clinical populations. It is possible that some clinical populations have more vigorous go-related activity vs. poorer stop-related activity during stopping. It could be used in developmental studies too. Fourth, our results are averaged across trials and participants, future studies could investigate the possibility of improving signal processing techniques such that these signals may be detected on single trials. Indeed, some studies have already demonstrated such techniques for event-related potentials (Hu et al., 2010). This would be of immense importance in the field as it would allow studying the trial-by-trial race between the go- and stop-process and help test several questions, for example, experimentally verify trigger failures.

Our results also speak to the debate over the existence of a ballistic stage. Over years, numerous studies have suggested the existence of a ballistic stage (Osman et al., 1986), while other have not (De Jong et al., 1990). While many of these studies have used simulations to validate the existence of a ballistic stage, our results provide neural evidence of the same. In other words, once the go-related activity reaches a certain threshold, muscles start to get activated and after a delay the response is initiated. A part of the final delay where the muscles are getting activated is non-ballistic where the stop-process can intervene and stop the response, consistent with numerous studies that have reported partial muscle responses which can be cancelled (Jana et al., 2020; Raud et al., 2022; Raud and Huster, 2017; Tatz et al., 2021). Here, we show that in the motor cortex, go-related activity can reduce ~100 ms prior to the response but still result in a response. That the response is not cancelled despite this ~100 ms window speaks in favor of the existence of a ballistic stage consistent with numerous studies (Gopal and Murthy, 2016; Jana et al., 2020; Jana and Murthy, 2018). It suggests that once the point-of-no-return is reached, a response is inevitable.

Recent studies have suggested that the stop-process is actually a two-stage process (Diesburg and Wessel, 2021; Schmidt et al., 2013; Schmidt and Berke, 2017; Tatz et al., 2021). A ‘pause’ mediated by the hyperdirect pathway which briefly raises the threshold for response, while the ‘cancel’ mediated by the indirect pathway which actually cancels the response. Our study is not able to discern between the putative two stages but it is possible that the beta activity reflects both the pause and cancel stages (Diesburg and Wessel, 2021). It is possible that the activity of slightly different beta frequency bands reflects the two processes. Future studies could investigate this.

While majority of our results pertain to action-stopping behavior, we also observed that prefrontal beta might be inherently inhibitory in nature. Notably, we observed beta activity above baseline in the go trials as well. This increased beta activity might relate to proactive slowing as the go trials were performed in the stop signal task which required one to stop in a minority of trials. Thus, beta activity might be of two types – intrinsic/proactive and reactive. Proactive beta may be detected when participants anticipate the necessity to stop or chose to slow down their responses, and reactive beta may be detected when participants rapidly need to stop their responses. Such proactive slowing in the go trials in the stop signal task has been reported previously (Soh et al., 2021). Proactive beta may have a retardive effect on go-related activity and in turn behavior, while reactive beta may result in inhibition of the response by increasing the degree of influence of the stop-on to the go-process. Whether or not these different types of beta originate from the same network needs to be tested further. Notably, such slowing of responses due to sensorimotor beta activity has been reported previously (Khanna and Carmena, 2017; Leriche et al., 2022; Little et al., 2019; Muralidharan et al., 2022; Pogosyan et al., 2009; Soh et al., 2021).

We also observed that the go-related activity in the slow RT trials reached a lower level compared to that in the fast RT trials. An interpretation of this is that intrinsic/proactive beta is retardive and lowers the decision threshold. This would seemingly be at odds with the recent suggestion that beta activity raises the threshold (Muralidharan et al., 2021). One possible solution of this conundrum is that beta may affect either the rate or threshold of accumulation. Another possibility is that beta affects the product of the rate and threshold, i.e. the integral of the accumulation. It is possible that for a response to be generated, the integral of the area under the go-related activity must reach a particular level and intrinsic beta might increases this level. In other words, beta might modulate the amount of integration that is required: trials where intrinsic beta is low, the decision variable has to integrate to a smaller level compared to trials where intrinsic beta is high. This level of integration could be changed both by modulating the threshold and/or the rate of accumulation. Thus, when raising the threshold but keeping the rate of accumulation constant, the amount of integration required would be higher compared to when the threshold is lower (a.k.a. (Muralidharan et al., 2022)). Similarly, lowering the threshold but substantially slowing the rate of accumulation could still increase the required level of integration for a response to be made (as observed here).

This study suffers from some limitations. Firstly, although the stop signal task was used in all the studies, they had subtle differences (e.g. duration of trials, number of practice trials, response buttons, etc.) which might explain some of the differences in results between the three data sets. Second, this study was motivated by the aim to discover neural correlates of stopping based on existing literature. Hence, we did not test all potential neural signatures or all possible methodological options. However, analyses were first carried out in one data set followed by replication in others. Third, the study is correlative. Future causal studies can test whether affecting the go-related and stop-related activity affects behavior.

In conclusion, this study demonstrates that neural correlates of the race model may be detected in scalp potentials and links the algorithmic and implemental level of our understanding of action-stopping. In the future, we envision that these neural correlates may be used to answer numerous extant questions regarding the neural and computational principles that govern action-stopping.

## Materials and methods

### Participants

Adult, healthy participants provided written informed consent and were compensated at $20/hour. The studies were approved by the Institutional Review Board of University of California San Diego (protocol #171285).

*Study 1*. Twenty-seven participants (sixteen females; age 21 ± 0.5 years; all but one right-handed). Five were excluded from analysis, two for technical failure with the EEG recording system and three as they lacked a right frontal brain IC based on our standard method (Castiglione et al., 2019; Wagner et al., 2018). This data set has been reported in a previous publication from the lab (Muralidharan et al., 2021).

*Study 2*. Fifteen participants (nine females; age 21 ± 0.4 years, all right-handed). Three were excluded from analysis, two for technical failure with the EEG recording system, while the other lacked a right frontal brain IC based on our standard method (Castiglione et al., 2019; Wagner et al., 2018). This data set has been reported in previous publications from the lab (Hannah et al., 2020; Jana et al., 2020; Muralidharan et al., 2021).

*Study 3*. Thirty-three participants (eighteen females; age 21 ± 0.3 years; all right-handed). Five were excluded from analysis as they lacked a right frontal brain IC based on our standard method (Castiglione et al., 2019; Wagner et al., 2018). This data set has been reported in a previous publication from the lab (Sundby et al., 2020).

### Stop signal task

This was run with MATLAB 2014b (Mathworks, USA) and Psychtoolbox (Brainard, 1997). Each trial began with a white square appearing at the center of the screen fixation cross. After 500 ± 50 ms, a right or left white arrow appeared at the center of the screen and the participant tried to press the corresponding left or right buttons with their right hand. The stimuli remained on the screen for 1 s. If participants did not respond within this time, the trial aborted, and ‘Too Slow’ was presented. On 25% of the trials, when the arrow turned red after a stop signal delay (SSD), participants tried to stop their response (Stop trials). The SSD was adjusted using two independent staircases (for right and left directions). SSDs increased and decreased by 50 ms following successful and failed stopping, respectively. Each trial was followed by an inter-trial interval (ITI) and the entire duration of each trial including the ITI was 2.5 s. Participants were requested to respond as quickly and as accurately as possible.

*Study 1*. Following 40 practice trials, participants performed 10 blocks of 40 trials each (300 go and 100 stop trials). To respond to the left and right arrows, participants had to press the left and right keys with their index and middle fingers, respectively.

*Study 2*. Following 80 practice trials, participants performed 24 blocks of 80 trials each (1440 go trials and 480 stop trials). To respond to the left arrow, participants had to press a key on a vertically oriented keypad using their index finger, while for the right arrow they had to press down on a key on a horizontally oriented keypad with their pinky finger.

*Study 3*. Following 40 practice trials, participants performed 4 blocks of 80 trials each (240 go and 80 stop trials). To respond to the left arrow, participants had to press a key on a vertically oriented keypad using their index finger, while for the right arrow they had to press down on a key on a horizontally oriented keypad with their pinky finger.

### EEG data recording

*Study 1*. 64 channel EEG data was recorded in the standard 10/20 configuration at 512 Hz using a ActiveTwo system (BioSemi Instrumentation, The Netherlands). External electrodes were placed on the bilateral mastoids and canthi, as well as below and above each eye. The data were on-line referenced to the BioSemi CMS-DRL electrode. Electrode offsets were maintained below 25 µV. *Study 2 & 3*. 64 channel EEG data was recorded in the standard 10/20 configuration at 1000 Hz using an actiChamps system (EasyCap, BrainVision LLC and Brain Products GMBH, Germany). Electrode Fpz served as the ground and additional electrodes were placed over each canthus and below the right eye. The data was recorded reference free and then offline re-referenced to electrodes placed bilaterally over mastoids. Electrode impedances were maintained <10 kΩ.

### EEG analysis

All data were analyzed using EEGLAB 14.1.1b (Delorme and Makeig, 2004) and custom scripts. For further details, please refer to the original publications as cited above.

*Study 1*. Data were downsampled to 512 Hz and high-pass filtered at 1 Hz (zero phase FIR filter, order 7500) to minimize slow drifts, and low pass filtered at 200 Hz (zero phase FIR filter, order 36). The EEG data were then re-referenced to a common average. Continuous data were visually inspected to remove bad channels and noisy stretches.

*Study 2*. Data were downsampled to 512 Hz and band-pass filtered between 2–100 Hz. A 60 Hz and 180 Hz FIR notch filter were applied to remove line noise and its harmonics. EEG data were then re-referenced to the average. Continuous data were visually inspected to remove bad channels and noisy stretches.

*Study 3*. Data were downsampled to 512 Hz and low-pass filtered with a MATLAB polyphaser antialiasing filter. We then re-referenced the data to an average of the two mastoid electrodes. We applied a high-pass filter at 2 Hz [finite impulse response (FIR) order 3,300] and a notch filter at 60 and 180 Hz (FIR order 846) to remove electrical noise. Continuous data were visually inspected to remove bad channels and noisy stretches.

After preprocessing, the noise-rejected data were then subjected to logistic Infomax Independent Component Analysis (ICA) to isolate ICs for each participant (Bell and Sejnowski, 1995). Using the DIPFIT toolbox in EEGLAB, we computed the best-fitting single equivalent dipole matched to the scalp projection for each IC (Delorme and Makeig, 2004; Oostenveld and Oostendorp, 2002). We identified and removed the activity related non-brain ICs related to eye movements, blinks muscle activity, and other sources using the frequency spectrum (increased power at high frequencies), scalp maps (activity outside the brain) and the residual variance of the dipole (greater than 15%). For each participant, a putative right frontal IC was then identified from the scalp maps (if not present then we used frontal topography) and whether there was increased beta activity between the stop signal and SSRT compared to fixation duration. The channel data were projected onto the corresponding right frontal IC.

### Selecting the stop-related activity

To circumvent potential ‘double-dipping’ confounds, we did not select the frequency band for each individual subject and then test the how its activity varied in the stop trials. Since the specific frequency can vary across participants, we also decided against choosing a fixed frequency band across all data set. Instead, we settled on a compromise where we could select a beta frequency band specific to each data set i.e., across all participants in each data. To do this, we averaged the mean power across all frequencies in the time window between the stop signal and SSRT in the successful stop trials. If there was a significant peak in the low beta frequency range (13-20 Hz), then we selected a window of ±3.5 Hz around this peak. If there was no noticeable peak then we selected the entire low beta range. For data sets 1 and 2, we did not detect a peak and hence selected the entire low beta range, while data set 3 we observed a peak at 15.25 Hz and hence selected a ±3.5 Hz window around this peak (chosen frequency in data set 1 = 13-20 Hz, data set 2 = 13-20 Hz, data set 3 = 11.75-18.75 Hz; ***Figure 2-figure supplement 1A-C***).

### Selecting the go-related activity

To circumvent potential ‘double-dipping’ confounds and also to be consistent with our selection of the stop-related activity, we did not select the frequency band for each specific subject and then test the how its activity varied in the go trials. Instead of choosing a fixed frequency range, we selected the frequency band specific to each data set, i.e., across all participants in each data set (the main results remained same even if we selected a fixed frequency range; data not shown). To do this, we averaged the mean power across all frequencies in the period between 300-600 ms after the go cue in the correct go trials as this is the time window where one observes a typical desynchronization in mu/alpha frequencies (Crone et al., 1998; Engel and Fries, 2010; Pfurtscheller and Lopes da Silva, 1999). We selected a window of ±3.5 Hz around the frequency which had the minimum power (chosen frequency range in data set 1 = 7.75-14.75 Hz, data set 2 = 7.25-14.25, data set 3 = 8-15 Hz; ***Figure 1-figure supplement 1A***). The period which was used to calculate the baseline activity was common for the stop-related and go-related activity – the mean frequency-specific activity across all trials during the baseline period of −400 to 0 ms before the go cue, i.e., the period when the fixation cue was present on screen.

### The relationship of the stop-related activity with SSD

To test the parametric relationship between stop-related activity and SSD we considered only specific SSDs. As the tasks used SSD staircasing, there was a wide range of SSDs. We selected the SSDs 100, 150, 200, and 250 ms were more represented across all participants. Further, we considered only those participants who had at least 10 trials in each of these SSDs. We then averaged the activity across all participants in a particular SSD and tested when, following the stop signal, was the activity significantly greater than zero (one-tailed test) for 10 consecutive ms (see statistical analysis section).

## Statistical analysis

To compare between time-series, we used the *statmod* function in EEGLAB using non-parametric permutation testing (‘perm’) and non-parametric bootstrap (‘bootstrap’) with 10000 iterations. In the figures, we depict the significance of the former and latter as a pink and grey bar, respectively. All tests were two-tailed unless specified otherwise. This was followed by false discovery rate correction using the MATLAB function *mafdr* using linear-step up procedure originally introduced by Benjamini and Hochberg (1995).

### Estimation of trigger failures using BEESTS

The percentage of trigger failures (pTF) was estimated using the Bayesian Estimation of Ex-Gaussian STop-Signal (BEESTS) model developed by Matzke and colleagues (Matzke et al., 2017). The BEESTS model presumes a race between two stochastically independent processes-a go- and a stop-process. The SSRT distribution is estimated using the go RT distribution, and by considering the failed stop RT as a censored go RT distribution. On every stop trial, the censoring points are sampled at random from the SSRT distribution. The RT distribution underlying the go-and stop-process is assumed to be ex-Gaussian with a gaussian and an exponential component and is characterized by 3 parameters (*μ*_*Go*_, *σ*_*Go*_, *τ*_*Go*_ and *μ*_*Stop*_, *σ*_*Stop*,_ *τ*_*Stop*_). The mean and variance of these distributions were determined as *μ* + *τ* and *σ*^2^ + *τ*^2^, respectively. The model uses Bayesian Parametric Estimation (BPE) to estimate the parameters of the distributions and pTF. A hierarchical BPE was used, where individual subject parameters are modeled with the group-level distributions as this approach is thought to be more accurate than fitting the data of individual participants and is effective when there is less data per participant (Matzke et al., 2013). The priors were bounded uniform distributions (*μ*_*Go*_, *μ*_*Stop*_: *U*(0,2); *σ*_*Go*_, *σ*_*Stop*_: *U*(0,0.5) *τ*_*Go*,_ *τ*_*Stop*_: *U*(0,0.5); pTF: U(0,1)). The posterior distributions were estimated using the Metropolis-within-Gibbs sampling and we ran multiple chains. We ran the model for 5000 samples with a thinning of 5. The Gelman-Rubin 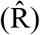 statistic was used to estimate the convergence of the chain. Chains were considered converged if 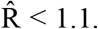 For further details about the model please refer to Heathcote et al. (2019).

## Data

The data is already publicly available, data set 1 at https://osf.io/3ersy/, data set 2 at https://osf.io/b2ng5/, and data set 3 at https://osf.io/6y5eb. On acceptance of the paper, the scripts used for this study will be uploaded on OSF.

## Acknowledgements

We thank Adam R. Aron for insightful comments on data analysis, Kelsey Sundby for sharing some EEG data, and Xinze Yu and Hunter Robbins for help in data recording. We gratefully acknowledge our support from Indian Institute of Technology Delhi, from National Institute of Mental Health (Grant No. MH-020002), and the James S. McDonnell Foundation (220020375).

