## Supplementary figures for "Bridging the neural and algorithmic correlates of action-stopping"

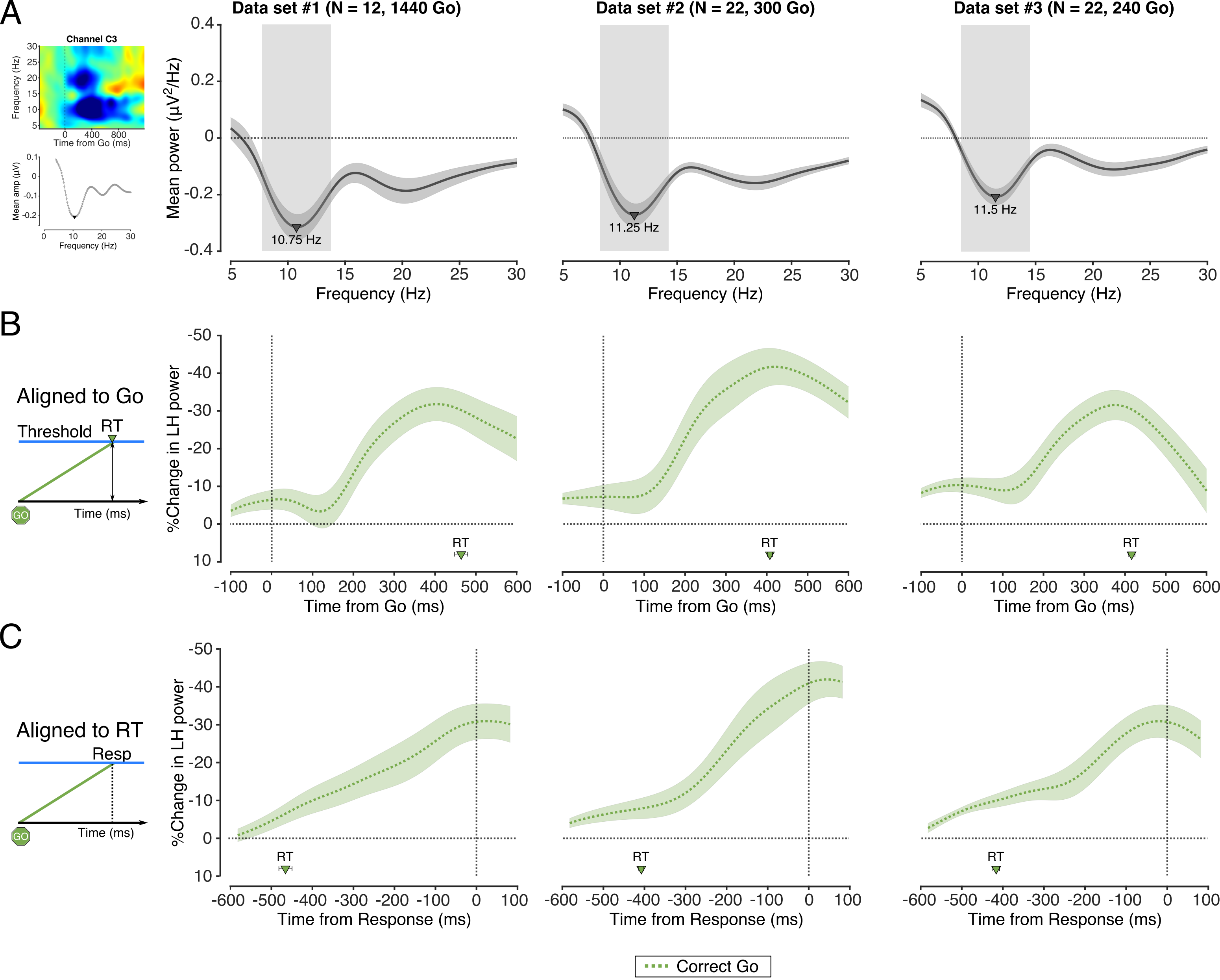


**Figure 1 - figure supplement 1 I Neural correlate of the go-process. A)** (*Left top*) The event-related spectral perturbation across all correct go trials in an exemplar participant time-locked to the time of the go cue. (*Left bottom*) The mean amplitude at all frequencies in that participant. (*Right*) The mean amplitude across all frequencies in all correct go trials across all participants in data set 1, 2, and 3. The shaded region represents the frequency range selected for the analyses - a ±3.5 Hz window around a central frequency (marked with a triangle). **B)** (*Left*) Schematic of the go-process rising to threshold, aligned to the go cue. (*Right*) The %change in LH power in the correct go trials across all participants, time-locked to the go cue, in data sets 1, 2, and 3. **C)** (*Left* ) Schematic of the go-process rising to threshold, aligned to the time of response, in the data sets 1, 2, and 3.

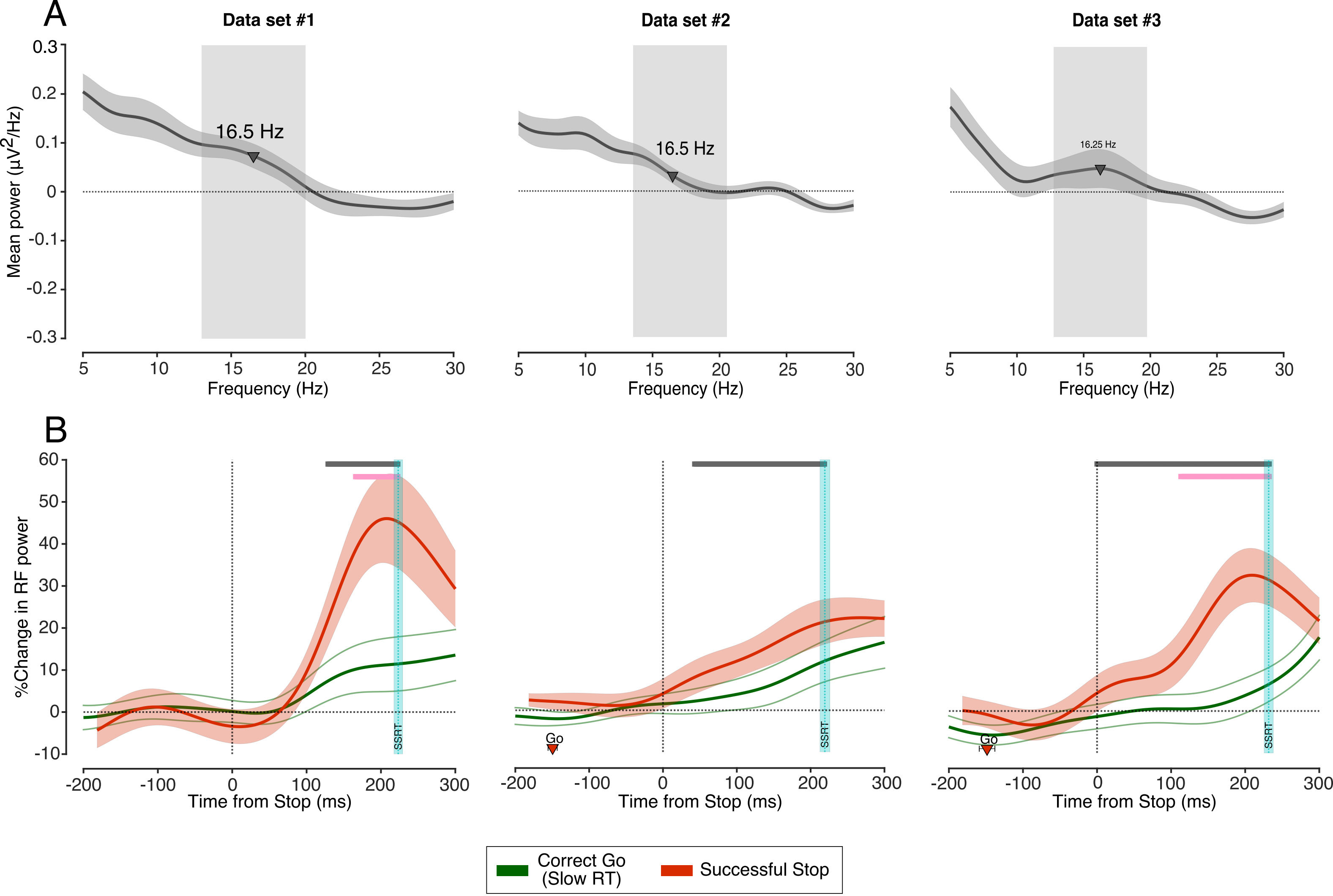


**Figure 2 - figure supplement 1 | Neural correlate of the stop-process**. **A)** Mean power in each frequency in the successful stop trials in the period between the stop signal and SSRT compared to the baseline, across all participants in a data set, for data set 1, 2, and 3. The shaded region represents the frequency range selected for the analyses, a ±3.5 Hz window centred around a specific frequency (marked with a triangle). **B)** The %change in the RF power aligned to the time of the stop signal compared between the successful stop and slower RT half of the correct go trials, in data sets 1, 2, and 3.

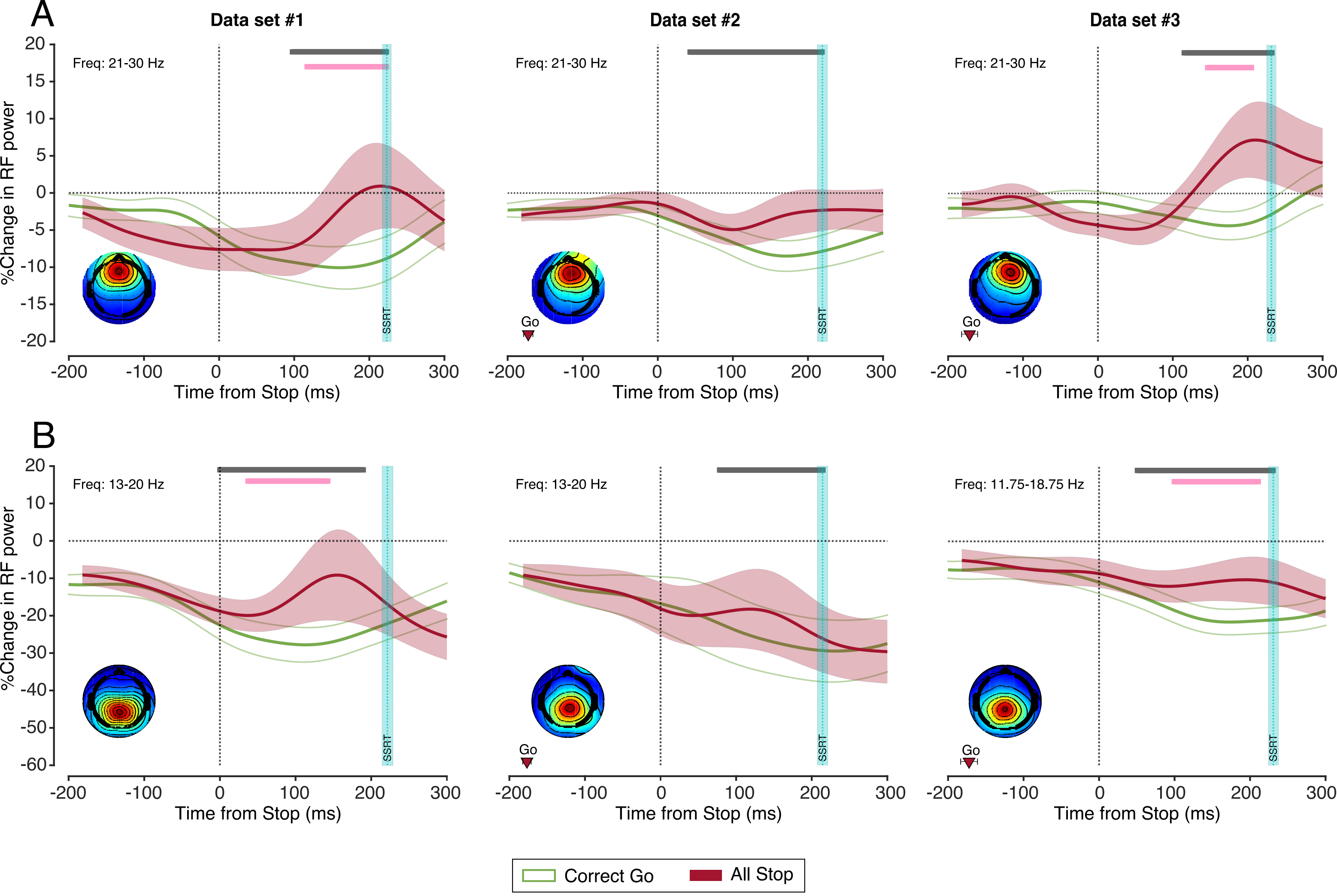


**Figure 2 - figure supplement 2 | Controls for the stop-related activity. A)** The %change in power in the 21-30 Hz frequency range aligned to the stop signal across all participants in all stop vs. correct go trials, in the data sets 1, 2, and 3. **B)** The %change in power in the RF frequency range aligned to the stop signal across all participants in all stop vs. correct go trials in the posterior IC, in the data sets 1, 2, and 3.
